# Convergent escape behavior from distinct visual processing of impending collision in fish and grasshoppers

**DOI:** 10.1101/2022.10.26.513903

**Authors:** Richard B. Dewell, Terri Carroll-Mikhail, Margaret R. Eisenbrandt, Thomas Preuss, Fabrizio Gabbiani

**Affiliations:** Department of Neuroscience, Baylor College of Medicine, Houston, Texas 77030; Hunter College and the Graduate Center, The City University of New York, New York 10065

**Keywords:** Looming, Collision-Avoidance, Neuron Models, Giant Fiber, LGMD, DCMD, Mauthner Cell

## Abstract

In animal species ranging from invertebrate to mammals, visually guided escape behaviors have been studied using looming stimuli, the two-dimensional expanding projection on a screen of an object approaching on a collision course at constant speed. The peak firing rate or membrane potential of neurons responding to looming stimuli often tracks a fixed threshold angular size of the approaching stimulus that contributes to the triggering of escape behaviors. To study whether this result holds more generally, we designed stimuli that simulate acceleration or deceleration over the course of object approach on a collision course. Under these conditions, we found that collision detecting neurons in grasshoppers were sensitive to acceleration whereas the triggering of escape behaviors was less so. In contrast, neurons in goldfish identified indirectly through the characteristic features of the escape behaviors they trigger, showed little sensitivity to acceleration. This closely mirrored a broader lack of sensitivity to acceleration of the goldfish escape behavior. Thus, although the sensory coding of simulated colliding stimuli with non-zero acceleration likely differs in grasshoppers and goldfish, the triggering of escape behaviors converges towards similar characteristics. Approaching stimuli with non-zero acceleration may help refine our understanding of neural computations underlying escape behaviors in a broad range of animal species.

**Key points:**

- A companion manuscript showed that two mathematical models of collision-detecting neurons in grasshoppers and goldfish make distinct predictions for their responses to simulated objects approaching on a collision course with non-zero acceleration.
- Testing these experimental predictions showed that grasshopper neurons are sensitive to acceleration while goldfish neurons are not, in agreement with the distinct models proposed previously in these species using constant velocity approaches.
- Both the escape behaviors of grasshopper and goldfish were insensitive to acceleration suggesting a further transformation downstream in grasshopper motor circuits that matches the computation observed in the goldfish Mauthner cell.
- Thus, in spite of different sensory processing in the two species, escape behaviors converge towards similar solutions.
- The use of object acceleration during approach on a collision course may help better understand the neural computations implemented for collision avoidance in a broad range of species.

## Introduction

Animals of different taxa are often preyed upon by the same or similar species. This raises the question of whether they have evolved convergent predator detection systems and avoidance behaviors. For visual detection of impending threats, diverse groups, including mammals, fish, insects, crustaceans, and birds possess neurons that specifically respond to visual looming stimuli, i.e., objects approaching on a collision course with constant velocity (Sun & Frost, 1998; Wu *et al*., 2005; Preuss *et al*., 2006; Fotowat *et al*., 2009; Nakagawa & Hongjian, 2010; Liu *et al*., 2011; de Vries & Clandinin, 2012; Oliva & Tomsic, 2014; Temizer *et al*., 2015; Dunn *et al*., 2016; Bhattacharyya *et al*., 2017; Bennett *et al*., 2019). These studies showed that the evoked behavior to visual looms differs between species, but response timing is comparable, typically occurring just before an impeding collision. This raises the question of how different (or similar) the neural computations that determine the timing of the behavior are.

Fish and grasshoppers react to visual looms, either with a powerful startle escape – triggered by a body bend (C-start) –, or with a prepared jump, propelling them away from the perceived threat (Fig. 1A, B; for C-starts in fish, see Preuss *et al*., 2006; Domenici & Hale, 2019; for jumps in grasshoppers, see Fotowat & Gabbiani, 2007; Fotowat *et al*., 2011). In fish, startle escapes to an overhead loom are typically initiated by a single action potential in one of the paired Mauthner cells (M-cells), which determines the timing and direction of the response (Fig. 1C). Loom stimuli presented from various directions, including from the front, or side also trigger escape behaviors in loosely restrained and freely moving zebrafish (Temizer *et al*., 2015; Bhattacharyya *et al*., 2017). Recordings from the M-cell ventral (visual) dendrite and chronic recordings from M-cell axons in freely behaving fish showed that peak M-cell membrane depolarization and action potentials in response to looms are tightly correlated and consistently occur just before collision time (Fig. 1 Cii, Ciii; Preuss *et al*., 2006; Weiss *et al*., 2006). In other words, the timing of C-starts measured behaviorally predicts the peak time of M-cell membrane depolarization.

**Figure 1.**
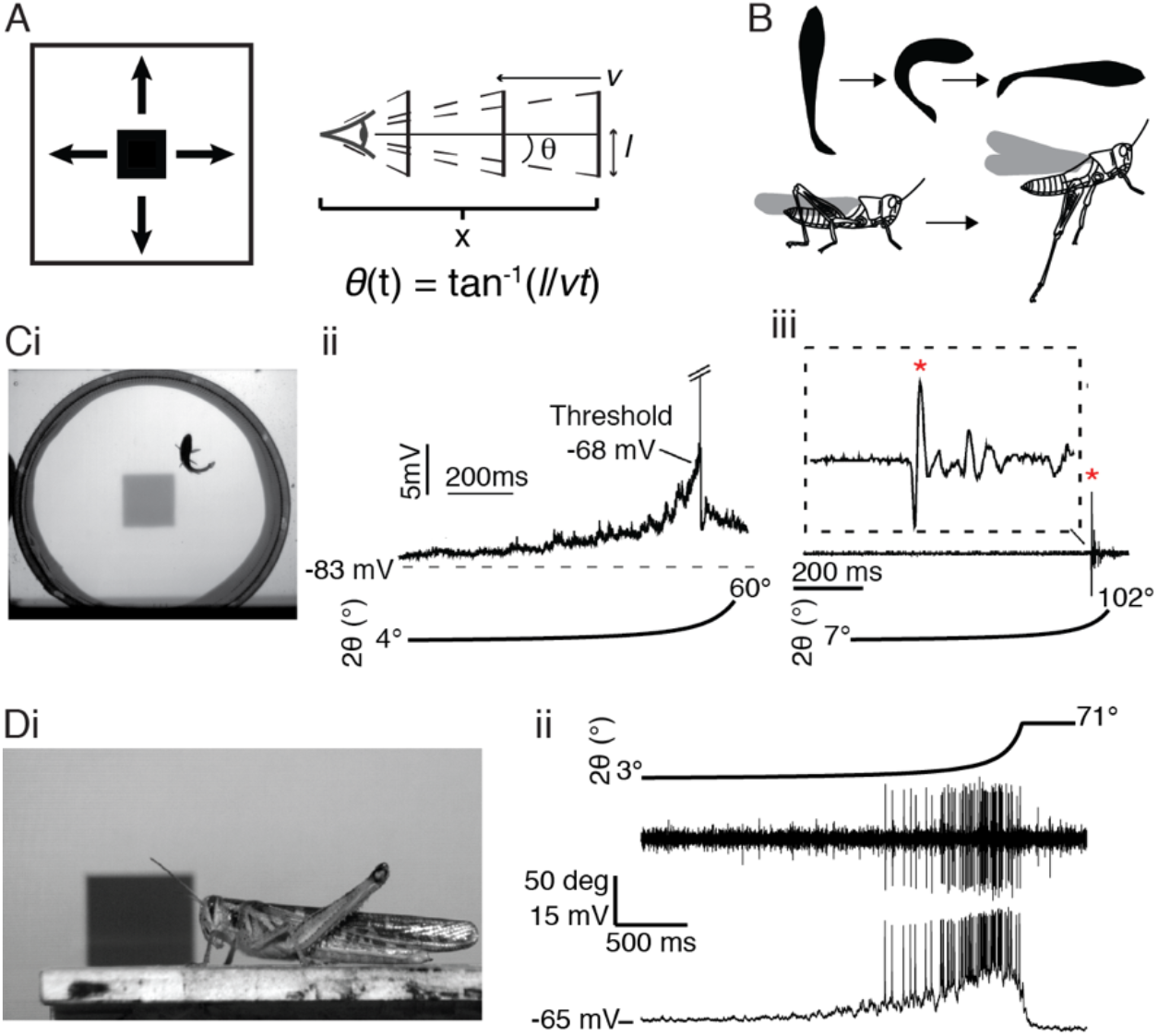
Fish and grasshoppers’ responses to looming stimuli. A) *Left*, illustration of a black looming stimulus, simulating a solid square with half-size *l* approaching with constant velocity *ν. Right*, the ratio *γ* = *l*/*ν* defines the angular half-size of the object *θ*(*t*) at time *t* relative to collision. B) Schematic illustrations of fish (top) and grasshopper (below) escape responses to looming stimuli. Ci) Ventral view of goldfish startle behavior (C-start) in response to a looming stimulus presented from above. Cii) Intracellular recorded M-cell membrane potential in response to a looming stimulus (stimulus angular size shown below). Ciii) Extracellular recording of M-axon field potentials in a freely behaving goldfish shows a single M-cell spike (*, magnified in dashed rectangular inset) preceding a loom-evoked C-start (not shown). Note that in both ii and iii the Mauthner cell spike occurs shortly before the end of loom expansion (modified from Preuss *et al*. 2006, and Weiss *et al*. 2006). Di) Image of a grasshopper being presented a looming stimulus from the side, taken before the animal jumped and flew away. Dii) Example simultaneous extracellular recording of DCMD spiking (middle) and intracellular LGMD membrane potential (bottom) during a looming stimulus presentation (top). The LGMD responds with high frequency firing that peaks shortly before the end of expansion.

In grasshoppers, escape behavior to simulated objects approaching from the side (Fig. 1Di) is triggered by the activation of an identified neuron called the descending contralateral movement detector (DCMD) which faithfully relays the firing pattern of the lobula giant movement detector (LGMD) neuron to motor circuits (O’Shea & Williams, 1974; O’Shea *et al*., 1974). In contrast to fish, the LGDM/DCMD neurons must generate sustained firing to trigger successful escape (Fig. 1Dii; Fotowat *et al*., 2011). This sustained firing (∼200 ms) likely contributes to escape jump preparation (Fotowat & Gabbiani, 2011). Yet, the firing rate of the DCMD neuron has a similar shape as the Mauthner cell’s membrane potential, peaking and decaying before projected collision time (Fig. 1Cii, Dii; Gabbiani *et al*., 1999).

Two distinct models have been proposed to describe the responses of goldfish and grasshoppers to simulated objects approaching at a constant speed towards the animal (Hatsopoulos *et al*., 1995; Gabbiani *et al*., 1999; Preuss *et al*., 2006). Here, we studied the neural responses predicted by these models to simulated accelerating approaching objects that have not yet been employed either in behavioral, or in electrophysiological experiments (see Gabbiani *et al.*, 2022). The same stimuli were used to compare and contrast the behavioral responses of fish and grasshoppers. We thus start by describing and comparing these novel accelerating stimuli to conventional constant speed looming stimuli. While the grasshopper and goldfish models make identical predictions for looming stimuli, they make distinct predictions for responses to accelerating stimuli, allowing to tease them apart and to better understand the differences in the neural computations implemented in these two model systems.

## Methods

### Animals

Grasshopper experiments were conducted with adult *Schistocerca americana*, 8-12 weeks of age. Animals were reared in a crowded laboratory colony under 12-hour light/dark conditions. Preference was given to large, active females ∼3 weeks after their final molt.

Fish experiments used nineteen naïve goldfish (*Carassius auratus*) of mixed sex, between 8.0 and 8.5 cm in standard body length, purchased from Ozark Fisheries, Stoutland, Missouri, USA. Fish were maintained in holding tanks (95 L; 30 × 30 × 60 cm; pH: 7.2-7.6, temperature: 19 ± 1 ºC), under 12-hour light/dark conditions and acclimated for at least two weeks prior to commencing behavioral experiments.

### Grasshopper behavioral setup

The stimuli were generated at 200 Hz with custom C code using the Scitech graphics library on a QNX4 PC. A cathode ray tube (CRT) monitor was used to present the stimuli. The monitor was positioned parallel to the animal’s body, centered on the eye to maximally stimulate it from the side at a distance of 6 cm. The animal was on a thin platform ensuring that the ipsilateral eye was between 5 and 7 cm from the stimulus. The luminance of the background was equal to 77 cd/m^2^, while dark simulated approaching objects had a luminance of 2 cd/m^2^. Behavioral responses were recorded with each video frame synched to the stimulus frames.

### Grasshopper behavioral experiment design

Grasshopper behavioral experiments were conducted as described previously (Fotowat & Gabbiani, 2007; Dewell & Gabbiani, 2018; Fig. 1B, D). Briefly, animals were placed on a short platform and walked along it parallel to the monitor. When the animal reached the end of the platform, placing their right eye in front of the center of the screen, a visual stimulus was started manually. The stimuli were presented in random order, and at least 10 minutes elapsed between trials of the same animal. In all, there were 739 trials using 24 animals and 9 stimuli (5 looms and 4 non-zero acceleration stimuli; see below for detailed description of the stimuli). Sufficient data were available from 11 animals (574 trials) for detailed within and across animal comparisons and these were used for subsequent analysis. For the experiments testing constant acceleration stimuli, 353 trials were recorded from 6 animals, with 10 trials excluded from analysis.

### Grasshopper behavioral data analysis

Jump probability was calculated based on the median unbiased estimator of a binomial response model. The jump timing (and corresponding subtended stimulus angle) were calculated based on the first video frame showing leg extension. Each video was examined to ensure the animal was properly positioned to see the stimulus, and trials in which the animal jumped or turned away before stimulus presentation were excluded. There weren’t enough trials to determine *α* and *δ* parameters of the *η* model for individual animals, so the *δ* used in threshold angle calculations was the same for all animals (see Results). The values were calculated by a weighted least squared-error linear fit of the jump time, *t*_*b*_, to looms as a function of the stimulus parameter, *γ* (Gabbiani *et al.*, 1999; Fotowat & Gabbiani, 2007; <u>see also *Grasshopper electrophysiological data analysis* below)</u>.

### Grasshopper electrophysiological experiments

For grasshopper electrophysiology, animals were placed ventral side up in a plastic holder with the left eye 18 cm from the stimulus monitor. The neck membrane was removed to expose the ventral nerve cord. Wires 50 μm in diameter were placed around the right nerve cord such that de-insulated wire was in contact with its dorso-medial surface allowing recording of action potentials from the descending contralateral movement detector (DCMD) axon coursing from the head to the thorax (Fig. 1D).

The same stimuli used in behavioral experiments were shown in randomized blocks with two minutes between stimuli. To prevent habituation, animals were exposed to regular visual, auditory, and mechano-sensory stimulation between trials. Responses to non-zero acceleration and looming stimuli were recorded from 11 animals. Responses to constant angular velocity and looming stimuli were recorded from 7 animals.

### Grasshopper electrophysiological data analysis

Extracellular recordings of DCMD activity were analyzed as previously described (Gabbiani *et al*., 1999; Dewell & Gabbiani, 2018). Briefly, DCMD action potentials were determined by amplitude thresholding and the instantaneous firing rate (IFR) was calculated by convolving the spike raster of each trial with a Gaussian filter having a standard deviation of 20 ms. The time of the peak response (*t*_*p*_) was measured as the time of the peak IFR relative to the projected time of stimulus collision. Normalized IFRs (Fig. 8G) were calculated by taking the mean IFR across animals in response to each stimulus and then dividing that by its peak value. This allowed to easily visualize the percentage of firing rate decay from the peak at the times of jump.

The *α* and *δ* parameters of the *η* model fits for looming stimuli were determined by a weighted least squared-error linear fit of the peak response time, *t*_*p*_, as a function of the stimulus parameter, *γ*. The parameter *δ* is the y-intercept of the fit, *α* is twice the slope (Gabbiani *et al*., 2022). The threshold angle was determined as the stimulus angular size at *t*_*p*_ − *δ* (Gabbiani *et al*., 1999). Fits of the *η* model for combined responses of looms and non-zero acceleration stimuli were determined using the Matlab least squared error minimization function ‘lsqcurvefit’.

### Fish experimental setup

Fish experiments were conducted in a tank measuring 77 cm in diameter and 30.5 cm in height that contained a round mesh arena insert of 39 cm in diameter and 27.6 cm in height. The tank was filled with conditioned water that matched that of the holding tanks. The water height was 7.5 cm, restricting the available depth of the swimming area. As a result, fish movement was mainly confined to a two-dimensional plane. A translucent white plastic lid on top of the tank served as projection screen for visual stimuli (screen-water surface distance 22.5 cm). The setup was situated on an anti-vibration table behind a curtain to avoid unintended mechano-sensory and visual stimuli.

Visual stimuli were projected on the screen with a digital light processing (DLP) projector (1024 × 768 pixels, 60 Hz; U4-131 Plus, Vision Corp., Tokyo, Japan). They were generated on a personal computer (PC) running the Windows operating system using Matlab and the Psychtoolbox (PTB-3). Overhead projection simulated predator approach relevant to *Carassius auratus* (Vijayan *et al*., 2018). At the screen, the luminance of the background was 2 cd/m^2^ and that of the visual stimulus simulating dark approaching objects was 18 cd/m^2^. To avoid habituation, inter-stimulus intervals were randomized between two and seven minutes. In addition, audio sound pips were delivered at random intervals for dishabituation (200 Hz; 5 ms; 158 dB rel. to 1 μPa in water). These stimuli were produced by a stimulator (Master-8, A.M.P.I., Jerusalem, Israel), amplified (Servo 120, Samson Technologies, Hickville, NY), and delivered via either one of two underwater loudspeakers (Model UW30, Electro-Voice, Burnsville, MN). To analyze the latency and kinematics of evoked startle escape behavior, a ventral view of the fish was recorded at 250 frame/s with a high-speed camera (AOS Technologies AG, Daettwil, Switzerland) and analyzed with imaging software (Imaging Studio, AOS Technologies AG, and ImageJ, NIH, Bethesda, MD).

### Fish behavioral experiment design

Individual fish were transferred from the holding tank to the experimental tank and acclimated for 15 minutes. In a first set of experiments, each animal (N=9) was exposed in 27 trials (±2) to nine different visual stimuli (∼3 trials per stimulus per subject). Stimuli were identical to those presented to grasshoppers, and likewise were presented in blocks with each stimulus shown once per block in random order. Stimulus angles were calculated based on the distance from the screen to the water surface. Five stimuli were looming stimuli with half-size (*l*) to speed (ν) ratios *l*/*ν* = −20, −30, −50, −70, and − 80 ms (Fig. 1A; see below for a detailed description of all visual stimuli). Two additional stimuli started their simulated approach as the looming stimulus with *l*/*ν* = −50 ms but accelerated to ‘collide’ at the same time as projected for looming stimuli with *l*/*ν* = −20, and − 30 ms, respectively. Similarly, two additional stimuli started their simulated approach as the *l*/*ν* = −50 ms stimulus but decelerated to collide at the same time as projected for looming stimuli with *l*/*ν* = −70, and − 80 ms. After an initial analysis of the evoked behavior, a second experiment (N=10 animals) focused on three looming stimuli with *l*/*ν* = −20, −50, and − 80 ms, and the corresponding accelerating and decelerating stimuli colliding at the same time as projected for those with *l*/*ν* = −20, and − 80 ms, respectively. This experiment had twice as many repeats of individual stimuli as the initial one (6, ±2). Slight differences in animal exposures to the stimuli were attributable to program errors or cessation of trials due to unresponsiveness of the subject. In total and across both experiments, 373 trials were conducted (75, ±4 for each of the stimuli with *l*/*ν* = −20, −50, and − 80 ms, as well as the associated *l*/*ν* = −20 accelerating, and *l*/*ν* = −80 ms decelerating stimuli).

All procedures were approved by and performed according to the Institutional Animal Care and Use Committee (IACUC) of Hunter College.

### Fish behavioral data analysis

Escape probability, latency, and C-start angular velocity were calculated based on frame-by-frame video analysis (frame rate: 250 Hz; Fig. 1B, C). Stimulus onset was determined as the first frame in which the stimulus was visible in the image. Latency was calculated as the time between stimulus onset and the first detectable head movement. Peak angular velocity of the initial part of the C-shaped body bend (Stage 1; Eaton *et al*., 2001) was measured in ImageJ as the change in angular direction of a line running from the fish’s center of mass to the tip of the head (see also Preuss *et al*., 2006; Weiss *et al*., 2006).

### Visual stimuli

We used three broad types of stimuli for behavioral and electrophysiological experiments in fish and grasshoppers: *(i)* looming stimuli with constant approach velocity; *(ii)* stimuli with constant, non-zero acceleration designed to ‘collide’ with the experimental subject at the same time as looming stimuli; and *(iii)* stimuli with constant angular speed of expansion. In all cases, the stimuli simulated, on a two-dimensional screen, an object approaching on a collision course towards the experimental subject. Yet, the three types of stimuli differ substantially in their approach trajectories, as detailed below. The first two types of stimuli were used to compare the timing of peak neuronal responses and of behavior to the predictions of the *η* and *κ* models previously used for grasshoppers and goldfish, respectively (see Results). The third stimulus type was required to further distinguish the two models for grasshopper electrophysiological and behavioral data. See Gabbiani *et al.*, 2022, for a detailed description of the stimuli and predicted responses.

Classically, looming stimuli (looms) have been defined as the two-dimensional simulation of objects approaching at a constant speed on a collision course with the animal (Schiff *et al*., 1962). For a solid square object of half-size *l*, starting at an initial distance *x*_*i*_ from the eye, the distance from the eye as a function of time *s* ≥ 0 measured from movement onset is described by

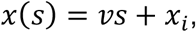

where *ν* < 0 is the approach speed. It will be useful to define a (dimensionless) normalized distance *y*(*s*) = *x*(*s*)/*l* in terms of the stimulus half-size so that

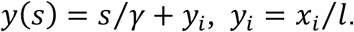

The constant *γ* = *l*/*ν* < 0 (in units of time) fully characterizes the approach trajectory as perceived at the animal’s eye (Gabbiani et al., 1999). In earlier work we have used the absolute value of *γ, l*/|*ν*| which provides an equivalent description, but the use of *γ* is more convenient in mathematical formulas. The time of collision *s*_*c*_ = −*γy*_*i*_ > 0 is obtained by solving the equation *y*(*s*) = 0. If time is referenced relative to collision, *t* = *s* − *s*_*c*_, then *y*(*t*) = *t*/*γ* and the half-angle subtended by the object at the retina is obtained by trigonometry as tan^−1^(*l*/*νt*) or

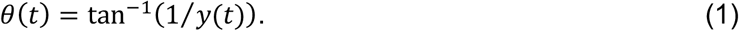

Five such stimuli are illustrated in Fig. 2A and B (green lines).

**Figure 2.**
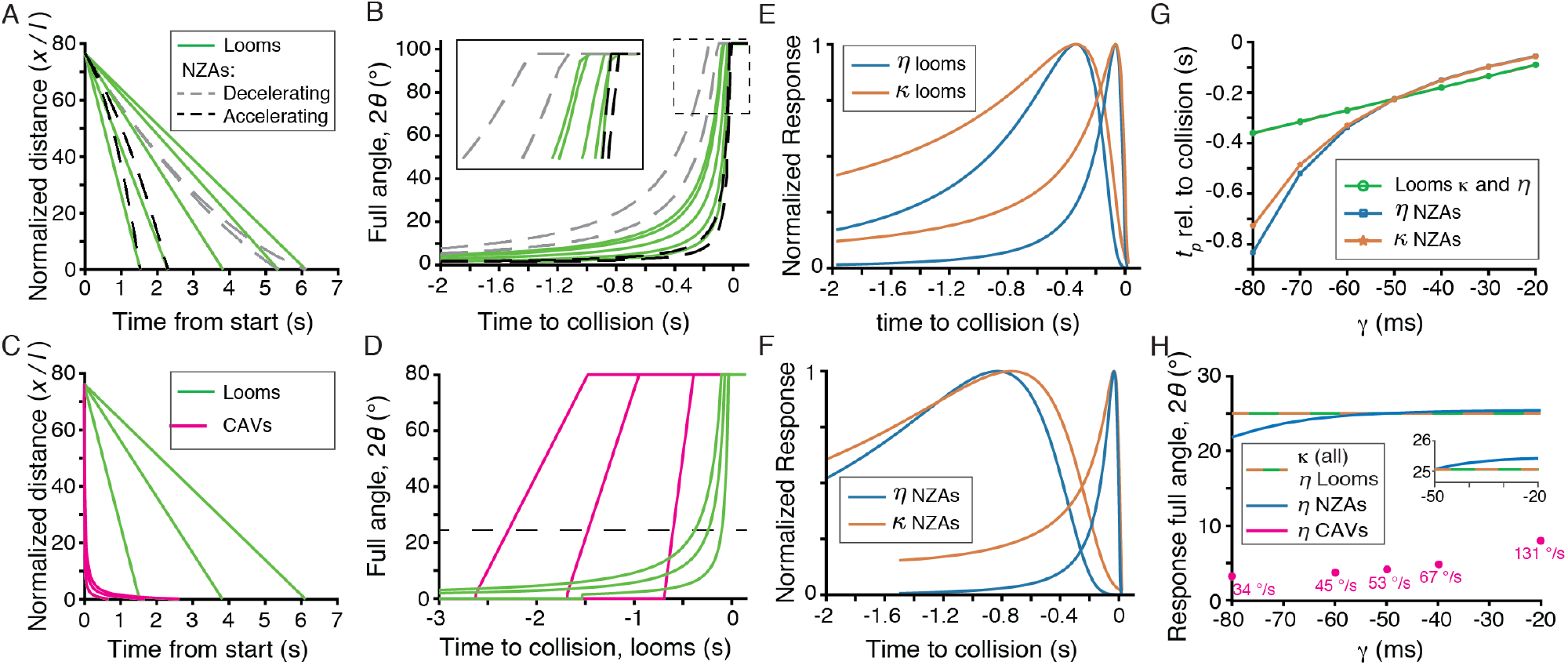
Visual stimuli simulating approaching objects with various velocity profiles and predicted model responses. A) Looming stimuli (looms, green) simulate objects approaching at constant velocity (i.e., a linear decrease in normalized distance); non-zero acceleration stimuli (NZAs) either accelerate (black) or decelerate (grey). B) Both looms and NZAs have steeply increasing angular sizes around the projected time of collision. The inset (solid box) magnifies angular trajectories close to collision time (dashed box in main panel). C) Compared to looms, constant angular velocity stimuli (CAVs; magenta lines) simulate faster initial approaches followed by steep deceleration. D) The angular size of CAVs increases linearly and their angular velocity (slope) was selected to match that of looms when 2*θ* equaled 25° (dashed line). E) Normalized time course of *η* and *κ* equations for looms with *γ* of −80 and −20 ms. For *η*, the plotted parameters were *α* = 9 and *δ* = 25 ms; for *κ*, the parameters were *β* = 4.573 and *δ* = 25 ms. F) Normalized time course of *η* and *κ* equations for NZAs with *γ* of −80 and −20 ms. G) The *η* and *κ* models predict identical peak response times to looms that are linearly dependent on *γ* (*P*green), but model-dependent response timings for NZAs (blue and ochre, respectively). H) Both the *η* and *κ* models predict a fixed peak response angle for looms. The *κ* model predicts the same peak response angle for NZAs and CAVs (dashed green-ochre line). The *η* model predicts slight changes for NZAs (blue line) and peak responses at smaller angles for CAVs (magenta dots, with the indicated constant angular velocity). CAVs are not parametrized by *γ*, hence for the *η* model, the predicted peak response full angles are plotted aligned to those looming stimulus *γ* values having matched angular velocity at their respective peak time (see Methods). The inset magnifies the plot for *γ* > −50 ms around the models’ pea) response angle (25°).

In our experiments, looming stimuli had an initial subtended half-angle of 0.75º (full angle: 1.5º) and expanded until filling the vertical axis of the screen. The maximum full angle (2*θ*) values reached by the stimuli were either 136º or 80º for the freely behaving or restrained grasshoppers, respectively, and 102º for the fish.

The normalized distance of stimuli with non-zero constant acceleration is given by

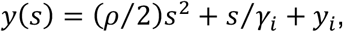

where *ρ* = *a*/*l* is the normalized accelerations in units of 1/time^2^ and 1/*γ*_*i*_ is the initial normalized speed (in units of 1/time). In our experiments, *γ*_*i*_ was always equal to −50 ms. Note that since *y*(*s*) > 0 decreases towards collision, *ρ* < 0 represents an accelerating stimulus (since the normalized distance decreases faster than for *ρ* = 0) and *ρ* > 0 a decelerating one (since the decrease in normalized distance will be slower than for *ρ* = 0).

Let us now consider a fixed looming stimulus with projected collision time *s*_*c*_ = −*γ*_*c*_*y*_*i*_. The above equation determines the normalized acceleration *ρ* required for a stimulus with initial normalized speed 1/*γ*_*i*_ to achieve the same projected collision time (as a function of *y*_*i*_, *γ*_*i*_ and *γ*_*c*_; see eq. 13 of Gabbiani *et al.*, 2022). In our experiments, we used two *γ*_*c*_ values to generate accelerating stimuli, *γ*_*c*_ = −20 ms so that *ρ* = −39.3 s^-2^ and *γ*_*c*_ = −30 ms so that *ρ* = −11.6 s^-2^. The two decelerating stimuli had *γ*_*c*_ = −70 ms so that *ρ* = 2.14 s^-2^ and *γ*_*c*_ = −80 ms so that *ρ* = 2.45 s^-2^. These non-zero acceleration stimuli (NZAs) are illustrated in Fig. 2A and B (dashed lines).

Constant angular velocity stimuli (CAVs) are defined by the equation

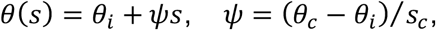

where *θ*_*i*_ is the initial angular size, *θ*_*c*_ = *π*/2 is the angular size at projected collision time (in radian), and *ψ* is the angular velocity in radian/time. If time is referenced relative to collision, this equation becomes

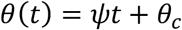

and the normalized distance is obtained from eq. (1), *y*(*t*) = 1/tan *θ*(*t*). From the three examples illustrated in Fig. 2C and D (pink lines), it can be seen that such stimuli represent a case of extreme deceleration compared to those considered above. Their normalized speed is largest at the beginning of approach and asymptotes at a value of −*ψ* at collision time (sec. 6 of Gabbiani *et al.*, 2022).

In grasshopper experiments, the angular velocities used for CAVs were selected to match the angular velocities of looming stimuli near the typical time of the grasshoppers’ peak neural response corresponding to a full threshold angle of 25º. For looms with *ψ* of −20, −40, −50, −60, and −80 ms this corresponds to *ψ* of 131, 67, 53, 45, and 34 º/s respectively (Fig. 2D, dashed line).

Although the term ‘looming stimuli’ has been used loosely recently, including for simulated constant angular velocity approaching stimuli, we will reserve our use of the terms ‘looming stimuli’ and ‘looms’ to the classical definition (constant approach speed). We will use the abbreviation NZAs for non-zero acceleration stimuli and CAVs for constant angular velocity stimuli.

### Statistics

All tests were two-tailed. Analyses of variance were calculated using Kruskal-Wallis non-parametric tests; the resulting significance levels are reported in the text as p_KW_. Significance of differences in response times or threshold angles between stimuli were calculated with the Wilcoxon rank sum test and reported as p_WRS_. For comparisons of behavioral response probabilities, significance was calculated using Fisher’s exact test and values are reported as p_FT_. Tests of difference from zero (Figure 7) were calculated using the Wilcoxon sign rank test and reported as p_WSR_. Comparison between fitted *η* model parameters for looms and CAVs used 10,000 bootstrapped estimates from their linear fits. The reported p-values for these comparisons, p_ASL_, were the ‘achieved significance level’ (ASL) statistic for two-sample testing of equality of means with unequal variance (Algorithm 16.2 in Efron & Tibshirani, 1993). The standard deviation is abbreviated by ‘sd’. The Pearson correlation coefficient is denoted by *ρ*_*P*_ and the p-value of the associated test for its difference from zero by p_*P*_.

## Results

### Stimuli simulating approaching threats differ widely in their characteristics

Classically, looming stimuli (or ‘looms’) have been defined as two-dimensional simulations on a screen of solid objects approaching on a collision course with the animal at constant speed (Schiff *et al*., 1962). In a plot of distance, *x*(*t*), from the eye as a function of time, they are represented by straight lines with slopes, *ν* < 0, since distance decreases as the object approaches. More relevant in this case is to plot the distance to the eye normalized by the half-size of the object as a function of time from movement onset, *y*(*t*) = *x*(*t*)/*l*, since it is directly related to the angular size of the stimulus (see eq. 1 and below; Fig. 2A, green lines). The inverse of the slope in this plot, *γ* = *l*/*ν* – or equivalently its absolute value, *l*/|*ν*| – may be used to characterize looming stimuli (in units of time; Gabbiani *et al*., 1999). Typical values triggering escape prior to collision in grasshoppers range from −20 to −120 ms (Fotowat & Gabbiani, 2007). In goldfish, prior experiments used values ranging from −10 to −50 ms to trigger escape (Preuss *et al*., 2006). The experiments described below used stimuli with *γ* = −80, −70, −50, −30 and − 20 ms for both grasshoppers and goldfish. Sensory stimulation on the animal’s retina is governed by the angular size subtended by the object, that may be computed from the normalized distance by trigonometry (Fig. 1A; eq. 1, Methods). For looming stimuli, the linear decline in distance results in a non-linear, nearly exponential increase in angular size as collision time nears (Fig. 2B, green lines).

In addition to these looming stimuli, we simulated objects that either accelerate or decelerate during approach (with the acceleration or deceleration, *a*, constant). We call these stimuli non-zero acceleration stimuli (NZAs). NZAs are characterized by their initial normalized distance *y*_*i*_, by their projected time of collision, and by their initial normalized speed, which in our experiments was always identical to that of a looming stimulus with *γ*_*i*_ = −50 ms. We selected the time of collision of looming stimuli with values *γ*_*c*_ = −80, −70, −30 and − 20 ms, allowing us to directly compare the behavioral and electrophysiological responses between NZAs and looming stimuli with matching collision times. Given *y*_*i*_, *γ*_*i*_ and *γ*_*c*_, one can determine algebraically the appropriate normalized acceleration, *ρ*(*y*_*i*_, *γ*_*i*_, *γ*_*c*_) (*ρ* = *a*/*l*, Methods). For *γ*_*c*_ < *γ*_*i*_ (i.e., *γ*_*c*_ = −80 and − 70 ms), the slope of the matching looming stimulus at collision time is shallower than the initial one (1/*γ*_*i*_), requiring deceleration (*ρ* = 2.45 and 2.14 1/s^2^, respectively). In this case, the trajectory is a portion of an upright parabola (Fig. 2A, grey dashed lines). Conversely, for *γ*_*c*_ > *γ*_*i*_ (i.e., *γ*_*c*_ = −30 and − 20 ms), the slope of the matching looming stimulus at collision time is steeper than that of the initial one, leading to acceleration (*ρ* = −11.6 and − 39.3 1/s^2^, respectively). In this case, the trajectory is a portion of an inverted parabola (Fig. 2A, black lines). Because at any given time before collision such a decelerating stimulus is closer than the looming stimulus with matching collision time, its angular size subtended at the eye is larger than that of the matching looming stimulus. The opposite holds for accelerating stimuli (Fig. 2B). Although NZAs have not yet been used experimentally, the normalized acceleration observed in prey capture attempts of vinegar flies by damselflies ranged from −2.14 to 0.71 s^-2^ (highest acceleration and lowest deceleration, respectively; sec. 5, Gabbiani *et al*., 2022; von Reyn *et al*., 2014). Hence our *ρ* values span a range encompassing what has been observed in these freely behaving insects.

The final set of stimuli used for behavioral and electrophysiological experiments in grasshoppers had a constant angular velocity (*ψ*; Fig. 2C, D). Hence, their angular half-size grows linearly from its initial value to reach 90º at collision time (Fig. 2D). The corresponding normalized distance reveals that they represent objects approaching rapidly initially, but then exhibiting strong time-dependent deceleration well ahead of collision time (Fig. 2C). We call such stimuli constant angular velocity stimuli (CAVs).

### Multiple models for neural responses to looming stimuli in different species

In grasshoppers, the LGMD/DCMD neurons are responsible for triggering jump escape behaviors to looming stimuli (Fotowat *et al*., 2011). The peak firing rate of these neurons comes a fixed neural delay after the looming stimulus reaches a threshold angular size, independent of the stimulus parameter *γ* (or equivalently, of its approach speed assuming a fixed half-size, *l*). A phenomenological model, called the *η* model, predicts this key aspect of their firing rate (Sun & Frost, 1998). In this model, the time-dependent firing rate is (up to a scaling factor) obtained by multiplying the angular speed of the object during approach by a non-linear function of its angular size:

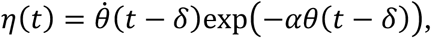

where *δ* is a neural delay between stimulus and response, *θ* is the half-angle (Fig. 1A), and 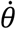 is the angular speed. The parameter *α* determines the threshold angular size as will become clear shortly. The *η* model predicts a firing rate that increases initially because 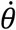 increases during the simulated approach. Yet, the object’s angular size also increases during approach. As a result, its negative exponential rapidly becomes vanishingly small, eventually turning off the *η* model’s response after it reaches a peak value (Fig. 2E). The timing of the firing rate peak is obtained by setting the time derivative of *η*(*t*) to zero, yielding the condition:

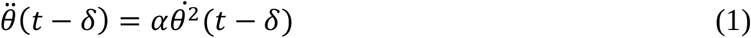

(sec. 4, Gabbiani *et al.*, 2022; Gabbiani *et al*., 1999). For looming stimuli, both 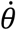 and 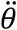 may be computed from the normalized distance *y*(*t*), leading to a linear relation between peak time, *t*_*p*_, and *γ*:

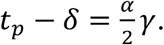

Because the normalized distance, *y*(*t*) = *t*/*γ*, is also a linear function of time with proportionality constant 1/*γ*, these two *γ* dependences cancel out when *t* = *t*_*p*_ − *δ* and the normalized distance is independent of *γ*, namely *y*(*t*_*p*_ − *δ*) = *α*/2. Equivalently, the half-angle at *t*_*p*_ − *δ* is given by

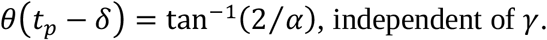

Various aspects of this phenomenological model have been tested experimentally (Gabbiani *et al*., 1999), including its possible mechanisms of biophysical implementation (Gabbiani *et al*., 2002; Jones & Gabbiani, 2010, 2012).

For goldfish a different model describing the membrane potential during looming was proposed, called the *κ* model (Preuss *et al*., 2006). In this model the membrane potential is (up to a scaling factor) a product of the stimulus angular size by a negative exponential of angular size:

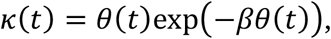

where we have omitted the neural delay *δ* for simplicity. Just as for the *η* model, this non-linear combination leads to an initial increase of *κ*(*t*) followed by a peak and an eventual decrease (Fig. 2E). The peak condition is obtained as for the *η* model,

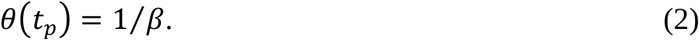

Thus, although the two models are different, they both predict a constant angular stimulus size at the peak time of their respective *η* and *κ* functions for looming stimuli (modulo a neural delay).

### The η and κ models predict distinct responses to NZAs

The *η* and *κ* models make identical predictions on the timing of peak responses for looming stimuli and are thus difficult to tease apart. But could their predictions diverge for other simulated approaching stimuli such as NZAs? That this is indeed the case may be seen by examining eqs. (1) and (2). The peak condition for the *κ* model (eq. 2) is directly constraining the angular half-size of the stimulus. Thus, peak responses always occur at a fixed threshold angular half-size, irrespective of stimulus type. In contrast, eq. (1) for the *η* model requires a specific relation between angular acceleration and speed during the simulated approach. This relation will not be fulfilled at the same time for NZAs and looming stimuli since the time dependence of their angular acceleration and angular speed differ (Fig. 2F).

The peak time of the *η* and *κ* models for NZAs can be computed analytically (sec. 5, Gabbiani et *al*., 2022). Assuming an angular half-size threshold of 12.5º as is typical for the DCMD peak firing rate, we find that both models predict earlier peak times for decelerating stimuli and later peak times for accelerating ones than for looming stimuli with the same *γ* value at collision time (Fig. 2G). For the *κ* model this can be explained by noting that the angular size of a decelerating stimulus is always larger than that of its matching looming stimulus (with parameter *γ*_*c*_; Fig. 2B). Thus, its peak angular threshold size must occur first. Conversely, the angular size of an accelerating stimulus is always smaller than that of the matching looming stimulus and thus its peak threshold angular size must occur closer to collision time (Fig. 2H). Hence, both the *η* and *κ* models predict a distinct arrangement of peak times for NZAs and looming stimuli that has not yet been tested experimentally.

Further, and as suggested by eqs. (1) and (2), the peak times in response to NZAs differ between the two models, with those of the *η* model occurring earlier for decelerating stimuli than those of the *κ* model and vice-versa for accelerating stimuli. The difference between the peak times predicted by the two models was, however, small for accelerating stimuli (∼1 ms; see Fig. 2G, *γ* > −50 ms). For decelerating stimuli, it increased with stronger deceleration (compare blue and ochre lines in Fig. 2G for *γ* < −50 ms). Thus the *η* model does not predict a constant angular threshold peak for NZAs, in contrast to the *κ* model (Fig. 2H). Analysis of surrogate data sets generated by the *η* and *κ* models using the previously described variability of DCMD responses (Gabbiani *et al*., 1999) suggested that the two models might be distinguishable experimentally (sec. 5, Gabbiani *et al.*, 2022).

### LGMD neuron’s responses to NZAs are consistent with the η model

To test whether the LGMD/DCMD neurons follow the predictions of the *η* or *κ* model, we recorded the DCMD spiking response to the looms and NZAs described above. Responses to decelerating stimuli were similar to looming responses, initially increasing and then decaying before collision time (Fig. 3A). Yet, the bulk of spikes occurred earlier than for their matched looms and so did the time of peak firing rate, as predicted by both models. The responses to accelerating stimuli also followed the general trend predicted by the models, with a tighter clustering of spikes and a peak firing rate slightly closer to collision time than for looms (Fig. 3B). For each animal, the peak times of responses to looms as a function of *γ* were fitted with a straight line, yielding a slope parameter, *α*, and a y-intercept, *δ* (Fig. 3C, green crosses and dashed line). The peak time responses to NZAs predicted by the *η* model based on these parameters yielded satisfactory fits as well (Fig. 3C, black crosses and grey dashed line). A fit of mean peak time responses for each animal and each *γ* value also yielded satisfactory fits and predicted well responses to NZAs (Fig. 3D). The range of values for the parameters *α* and *δ* obtained from these experiments agreed with those found in previous studies (Fig. 3E; e.g., Gabbiani *et al*., 1999). The angular threshold size computed from these fits were not significantly different from one another for looms (Fig. 3F). In contrast, NZAs yielded significantly different values for accelerating and decelerating NZAs, consistent with the *η* model but not with the *κ* model.

**Figure 3.**
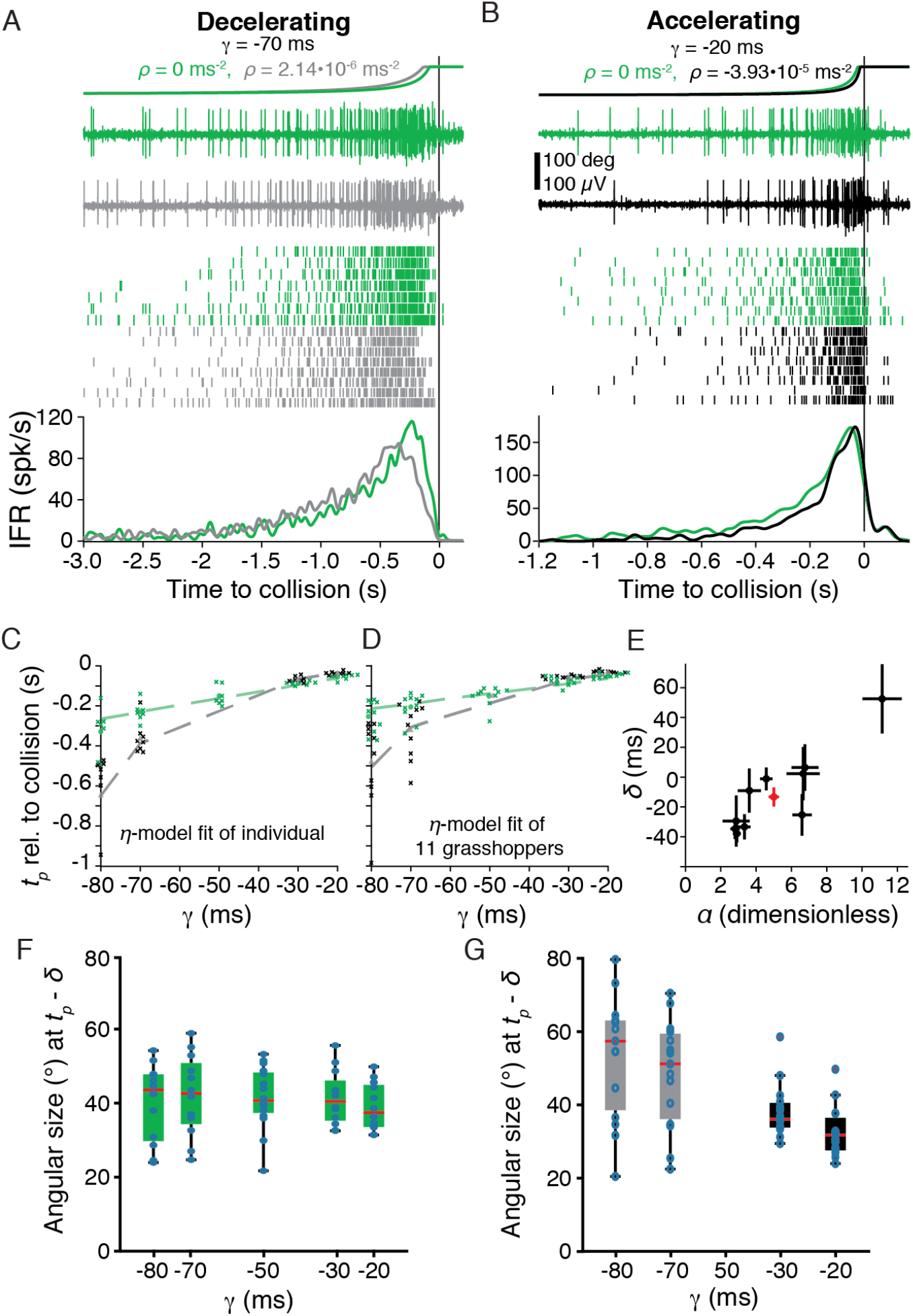
Acceleration shifts the timing of grasshoppers’ neural responses. A) *Top*, the angles of simulated objects approaching at constant velocity (looms, green) and decreasing approach velocity (decelerating NZAs, grey). *Immediately below, from top to bottom*, example nerve cord recordings, spike rasters, and plot of average instantaneous firing rate (IFR) show DCMD responses to the two stimuli. The peak response time was earlier for decelerating NZAs. B) Responses to stimuli with increasing approach velocity (accelerating NZAs, black) were delayed relative to constant velocity stimuli with the same *γ* (looms, green). Data from the same animal as shown in A. C) Peak DCMD response times of each trial (crosses) and the best fit *η* function for the animal shown in A and B (dashed lines). D) Fit of the population data; crosses are the mean response of each animal. In C and D, data points have been shifted horizontally to improve visibility. Green discs and grey triangles are mean of data points. E) The best fit *α* and *δ* parameters ± their bootstrapped standard errors for each grasshopper (black) and the population (red). F) The threshold angular size preceding the peak DCMD response for each stimulus. Constant velocity stimulus responses had angular thresholds independent of the stimulus parameter *γ* (p_KW_ = 0.96). G) Response angle changed with acceleration (p_KW_ = 0.01). (For D-F, N = 11 animals.)

### Grasshopper jump escape behavior differs for NZAs and looms

We next compared the jump escape behavior of grasshoppers to looms and NZAs. In these experiments, grasshoppers jumped in response to 58% of looms, 58% of decelerating NZAs and 50% of accelerating NZAs (Fig. 4A). No difference was detected among these three values (p_KW_ = 0.61) and the last two jump probabilities were not significantly different either (p_FT_ = 0.21). As predicted by the *η* and *κ* models, the jump times occurred earlier for decelerating, and later for accelerating NZAs than for their matched looms (Fig. 4B). Consistent with observations made on looms over the same range of *γ* values (Gabbiani *et al*., 1999; Fotowat & Gabbiani, 2007), the jump time variability was higher for decelerating than accelerating NZAs. The observed differences reached statistical significance only for the largest deceleration and acceleration values. As reported in freely behaving flies, the probability of escape before collision was smaller for accelerating than decelerating stimuli (Fig. 4C; von Reyn et al. 2014). It was also larger for stimuli with lower approach velocities than for stimuli with higher approach velocities (smaller, resp. larger *γ* values for fixed *l*; Fig. 4C). The fraction of ‘successful’ escape jumps, defined as those occurring before collision, was also higher for decelerating than accelerating NZAs, as well as for stimuli with lower approach velocities (smaller *γ* values, Fig. 4D). Fitting a straight line to the escape behavior time relative to collision as a function of *γ* yielded an estimated slope, *α*, and a y-intercept, *δ*, as for the time of peak DCMD response. The associated threshold angle at a delay *δ* prior to escape was more variable than that of peak DCMD firing and not significantly different across looms (Fig. 4E; compare with Fig. 3F). In contrast to the DCMD peak firing threshold angle, no significant differences were observed in escape threshold angles for NZAs (Fig. 4F). Thus, grasshopper escape behavior follows more closely the *κ* than the *η* model.

**Figure 4.**
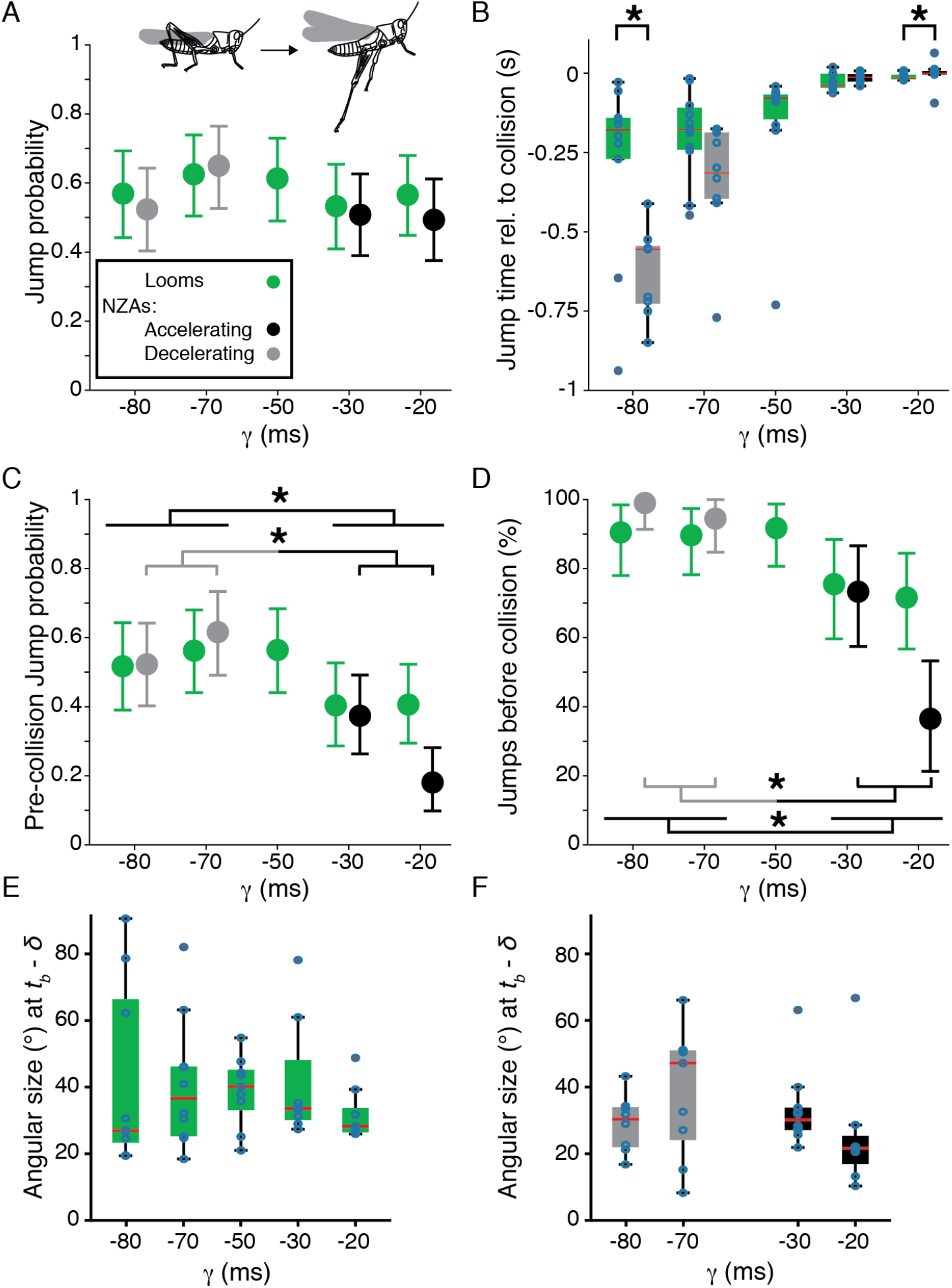
Grasshopper escape timing changes with stimulus acceleration. A) Grasshoppers jumped in response to all stimuli with similar probability (p_KW_ = 0.61). B) Stimulus acceleration changed the response time. Acceleration produced later jumps and deceleration produced earlier jumps (*: p_WRS_ < 0.05). C) The probability of producing jumps before the time of collision differs across stimuli (p_KW_ = 0.007) with fewer jumps before collision for accelerating stimuli than decelerating ones (*: p_FT_ = 2.6•10^−6^) and for faster stimuli (*γ* > −50 ms) versus slower ones (*γ* < −50 ms; *: p_FT_ = 1.3•10^−6^). D) Faster stimuli similarly produced a lower percentage of jumps that occurred before the time of collision; decelerating stimuli elicited higher jump percentages before collision than accelerating ones (*: p_FT_ = 1.4•10^−9^) and percentages for *γ* < −50 ms were higher than for *γ* > −50 ms (*: p_FT_ = 4.6•10^−10^). E) There was no difference in response threshold angle for looms (p_KW_ = 0.69; *δ* = 57 ms). F) Response threshold angle was not different among NZAs (p_KW_ = 0.18). N=11 animals for all plots. *t*_*b*_: time of escape behavior.

### Fish escape behavior to NZAs is more consistent with the κ than the η model

Similar to grasshoppers, the probability of escape to looms and NZAs was comparable in goldfish with responses to 69% of looms, 72% of accelerating NZAs, and 73% of decelerating NZAs (Fig. 5A). In contrast to the predictions of the *η* and *κ* model, we observed no significant difference in escape timing when comparing accelerating or decelerating NZAs with looms (Fig. 5B). We used the same linear fitting procedure applied to grasshopper peak DCMD firing and jump escape times to estimate the threshold angle triggering escape, and the delay between this angular threshold and behavior. No significant differences between threshold angles to looms and NZAs were found (Fig. 5C).

**Figure 5.**
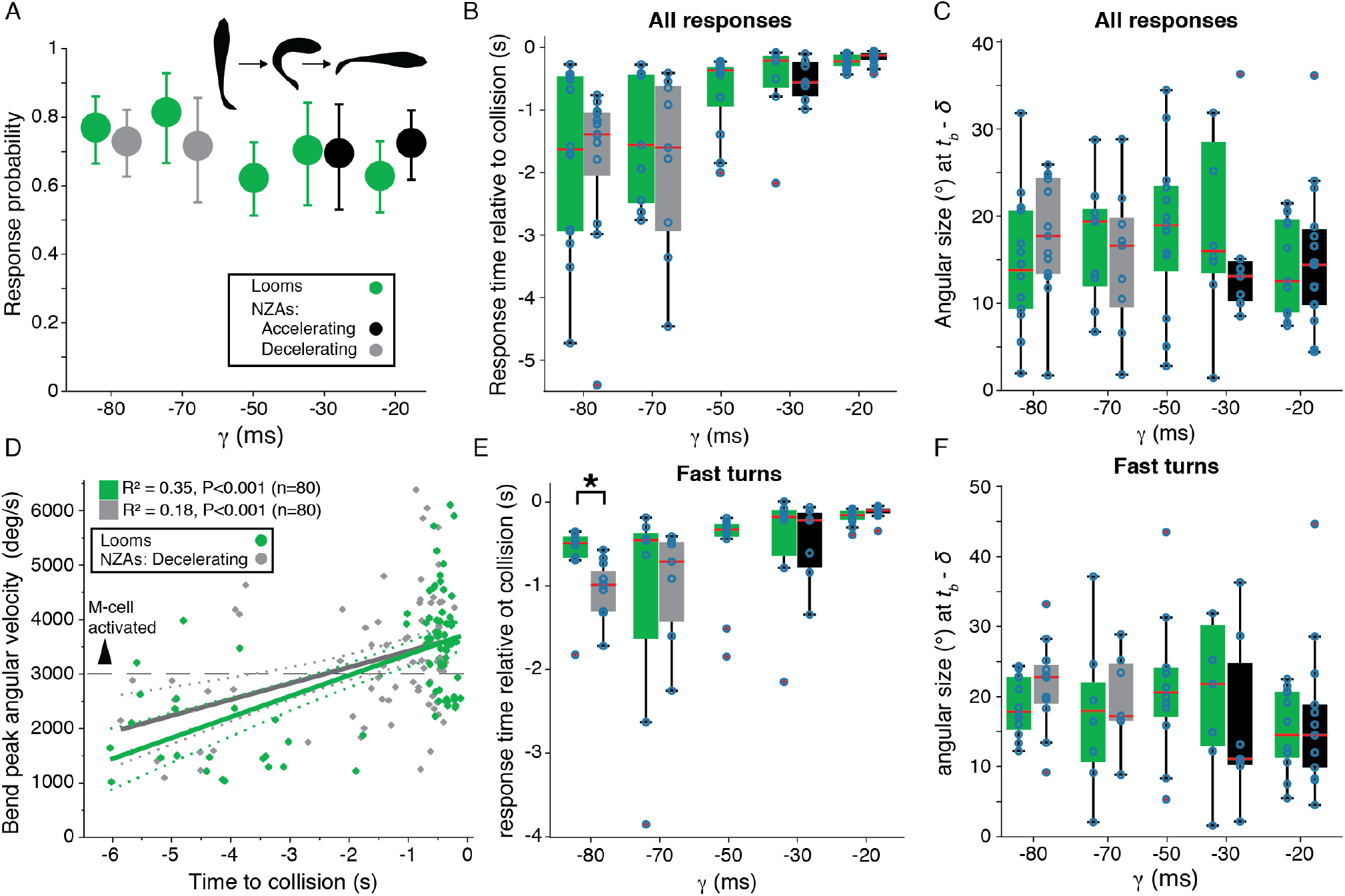
Fish escapes (C-starts) are not affected by stimulus acceleration. A) C-start (inset) probability was comparable for all loom approach velocities (p_KW_ = 0.38). B) Response timing changed with *γ* for both looms and NZAs (p_KW_ < 0.001), but the NZAs escape timings did not differ from their paired looms (p_WRS_ ≥ 0.16). C) The response threshold angle (*δ* = 25 ms) was comparable across stimuli (p_KW_ = 0.67) without significant differences between looms and NZAs with equal *γ* values (p_WRS_ ≥ 0.15). D) The peak angular velocity of C-starts increased for responses occurring closer to collision time. Dashed horizontal line indicates C-starts with velocity over 3000°/s which typically are initiated by the Mauthner neuron (see Fig. 1C). E). The timing of fast turns in response to NZAs differed from looms with equal *γ* only for the largest deceleration (*γ* = −80 ms; *: p_WRS_ = 0.002). F) The threshold angle for fast turns was not different across looms (p_KW_ = 0.38) or NZAs (p_KW_ = 0.18). N=19 animals for all plots.

The C-start escape turns triggered by the Mauthner cell usually result in peak angular turn speeds in excess of 3000 º/s (Eaton *et al*., 1981; Preuss *et al*., 2006; Weiss *et al*., 2006). For looms and decelerating NZAs, we found a positive correlation between escape time relative to collision and angular turn speed (Fig. 5D). A similar analysis could not be carried out for accelerating stimuli as the triggered escape behaviors clustered close to projected collision and were thus not sufficiently spread out in time to assess correlation between these two variables. Indeed, across stimuli, the fast bends occurred later than the slow bends (p_WRS_ = 3.7•10^−7^). Only 18% of early responses (>1.75 s before collision) were fast bends, whereas 70% of them were in the last 1.75 s before collision. The fast bend escape timing of NZAs and looms were more similar to the predictions of *η* and *κ* models, with decelerating NZAs responses occurring earlier than those of looms (Fig. 5E). There were no differences in the threshold angles for the fast turn responses to looms or NZAs (Fig. 5F). This constant threshold angle independent of acceleration matches the prediction of the *κ* model.

### Fish respond to faster stimulus expansion with more fast bends

Since most fast bend escape responses occurred closer to the time of collision, this suggested that a larger fraction of fast, Mauthner-mediated C-starts might occur for faster approaching stimuli (larger *γ* values). To confirm this hypothesis, we examined the probability of slow and fast bend responses for looms and NZAs. Slower expanding stimuli (*γ* < -50 ms) produced more slow turn responses (Fig. 6A). Similarly, decelerating stimuli produced more slow turns than did accelerating ones (Fig. 6A). Conversely, faster stimuli and accelerating ones produced a higher probability of fast turns (Fig. 6B). The majority of escape responses recorded in our experiments had peak bend velocities over 3000 º/s, indicating that they are likely initiated by the Mauthner neuron. The fraction of putative Mauthner initiated escapes increased with stimulus speed (i.e., for larger *γ* values; Fig. 6C). A modest positive linear trend was found between fast turn fraction and the stimulus parameter *γ* (*ρ*_*P*_ = 0.26, p_*P*_ = 5.8•10^−7^). These data suggest that faster or accelerating predator approaches are more likely to activate the Mauthner cell, in agreement with zebrafish results and similar to the increase in GF activation observed in flies for these stimuli (von Reyn *et al*., 2014; Bhattacharyya *et al*., 2017).

**Figure 6.**
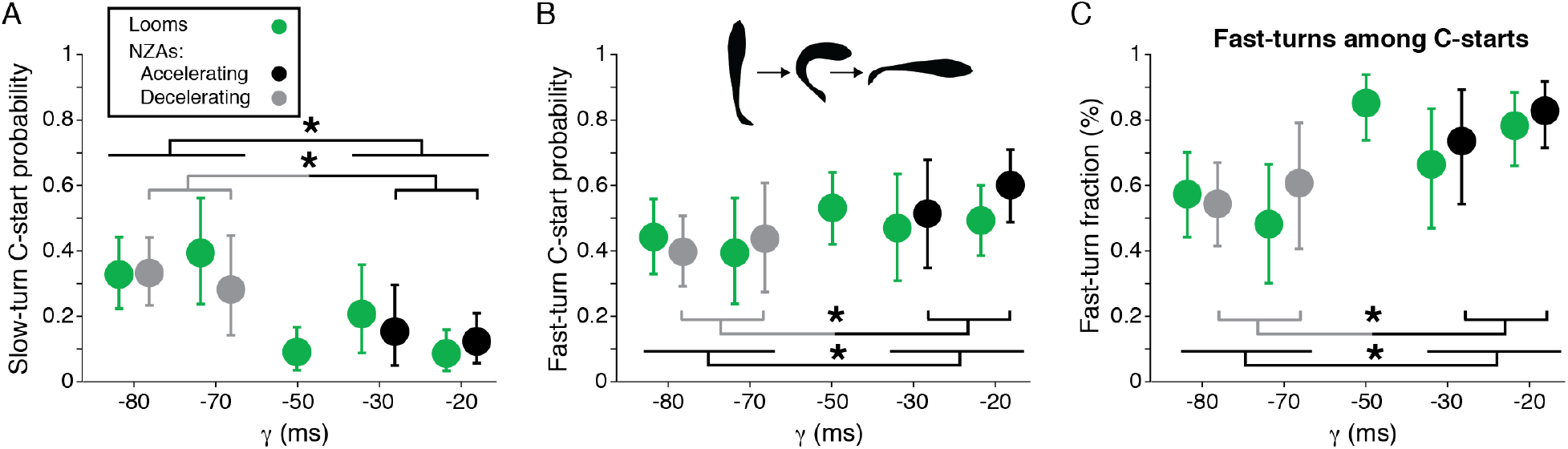
Faster simulated approaches (larger *γ* values) produce more fast-turn responses in goldfish. A) The probability of slow-turn C-starts was higher for slower simulated approaches; the slow-turn probability for *γ* < −50 ms was higher than for *γ* > −50 ms (*: p_FT_ = 1.1•10^−5^) and the slow-turn probability for accelerating stimuli was lower than for decelerating ones (*: p_FT_ = 0.002). B) Conversely, the probability of fast-turn C-starts was higher for faster simulated approaches; it was higher for accelerating than decelerating stimuli (*: p_FT_ = 0.014) and it was also higher for *γ* > −50 ms than for *γ* < −50 ms (*: p_FT_ = 0.005). C) The fraction of fast-turns among all C-start responses also increased as the simulated approach velocity increased; it was higher for accelerating than decelerating stimuli (*: p_FT_ = 0.0009) and it was higher for *γ* > −50 ms than for *γ* < −50 ms (*: p_FT_ = 2.0•10^−6^).

### Within-animal comparisons of response timings further refine population results

For each grasshopper and goldfish, we examined the within-animal change in response timing between looms and NZAs of equal *γ*. For neural responses in grasshoppers, acceleration delayed peak firing and deceleration produced earlier peaks, changes consistent with both the *η* and *κ* models (Fig. 7A, left). The influence of acceleration on the timing of grasshopper behavior was similar to that of the neural response, but the changes weren’t significant for the decelerating stimuli (Fig. 7A, center). The goldfish behavior showed no consistent change in response timing with acceleration, inconsistent with either model prediction (Fig. 7A, right).

**Figure 7.**
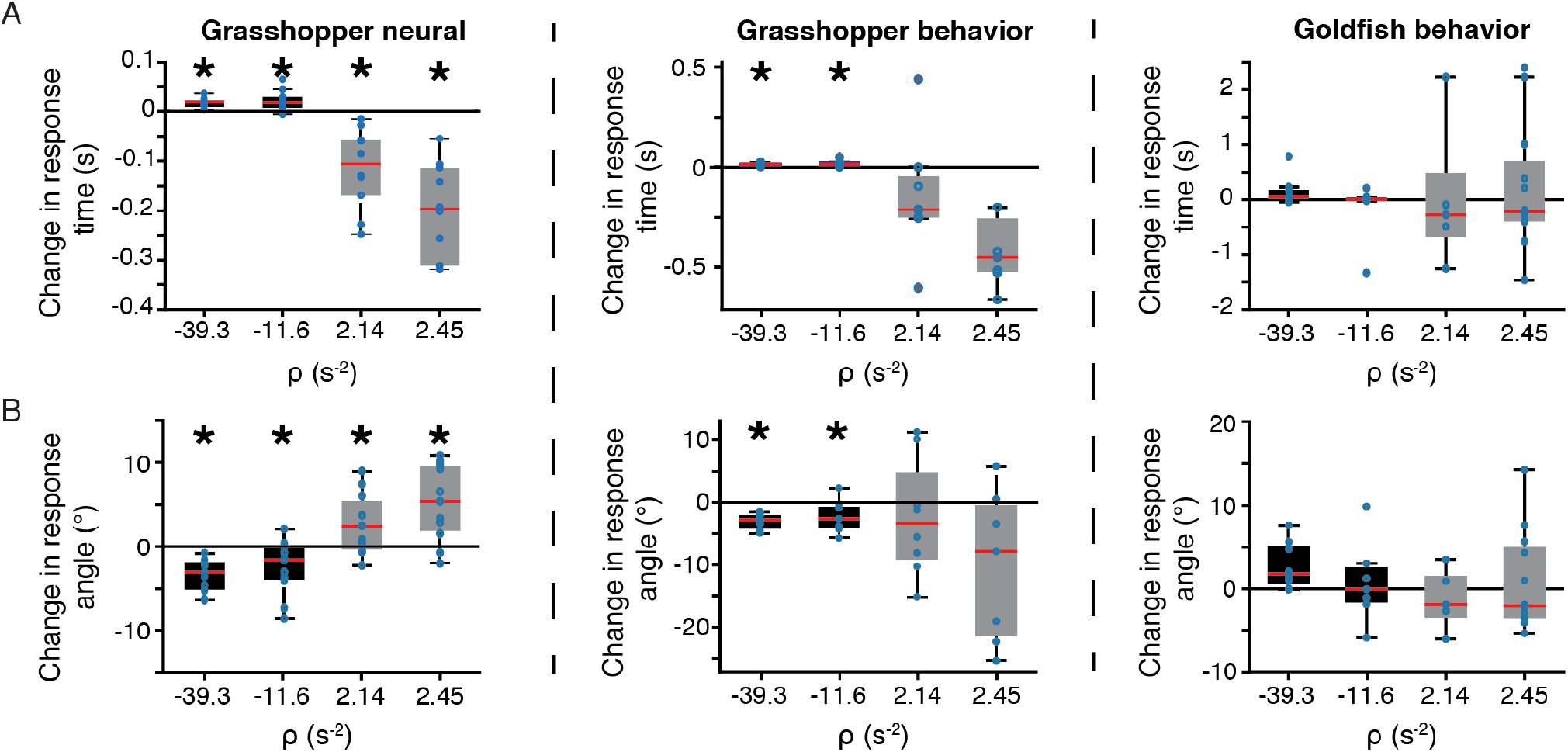
Comparison between grasshopper and fish responses. A) Within animal comparisons showed that grasshoppers responded later to accelerating stimuli (black boxes) and earlier to decelerating stimuli (grey boxes) whereas no timing difference was found in fish. B) Within animal comparisons of the threshold angle of grasshopper neural responses showed an increase with deceleration relative to looms. In contrast, the threshold angle at behavioral onset was not affected by acceleration in either species. *: p_WSR_ < 0.05.

The threshold angle of the grasshopper neural response was changed by acceleration (Fig. 7B, left). This change is inconsistent with the *κ* model, but, unexpectedly, the direction of the change was not that predicted by the *η* model (c.f. Fig. 2H). For grasshopper behavior, the median threshold angle was smaller for the two accelerating stimuli, matching the neural timing (Fig. 7B, center). The goldfish response threshold angle was unchanged by stimulus acceleration, consistent with the *κ* but not with the *η* model prediction (Fig. 7B, right).

### CAVs elicit different timings for neural and behavioral responses in grasshoppers

The grasshopper neural responses to NZAs showed changes in timing that were acceleration-dependent, and more consistent with the *η* than the *κ* model. The timing of the grasshopper behavioral response was inconclusive for distinguishing the models. We thus used constant angular velocity stimuli (CAVs), as the *η* and *κ* models predict large timing differences for them (Fig. 2H). Specifically, the *η* model predicts a peak response to CAVs a fixed delay (*δ*) after the stimulus begins expanding, while the *κ* model predicts a peak response at a delay *δ* after a threshold angular size, and hence a delay from stimulation onset that decreases with increasing angular velocity (sec. 6, Gabbiani *et al.*, 2022).

Looms produced the characteristic ramp up and decrease in firing rate as the stimulus expanded (Fig. 8A). In contrast, CAVs produced firing patterns that increased quickly to a peak and then decayed more slowly (Fig. 8B). The time of peak responses to CAVs was independent of the expansion speed unlike for looms (Fig. 8C). The threshold angle was fixed for looms but changed with the angular speed of CAVs (Fig. 8D). The timing and angular thresholds extracted from these data confirm that the *η* model best describes the grasshopper neural response.

**Figure 8.**
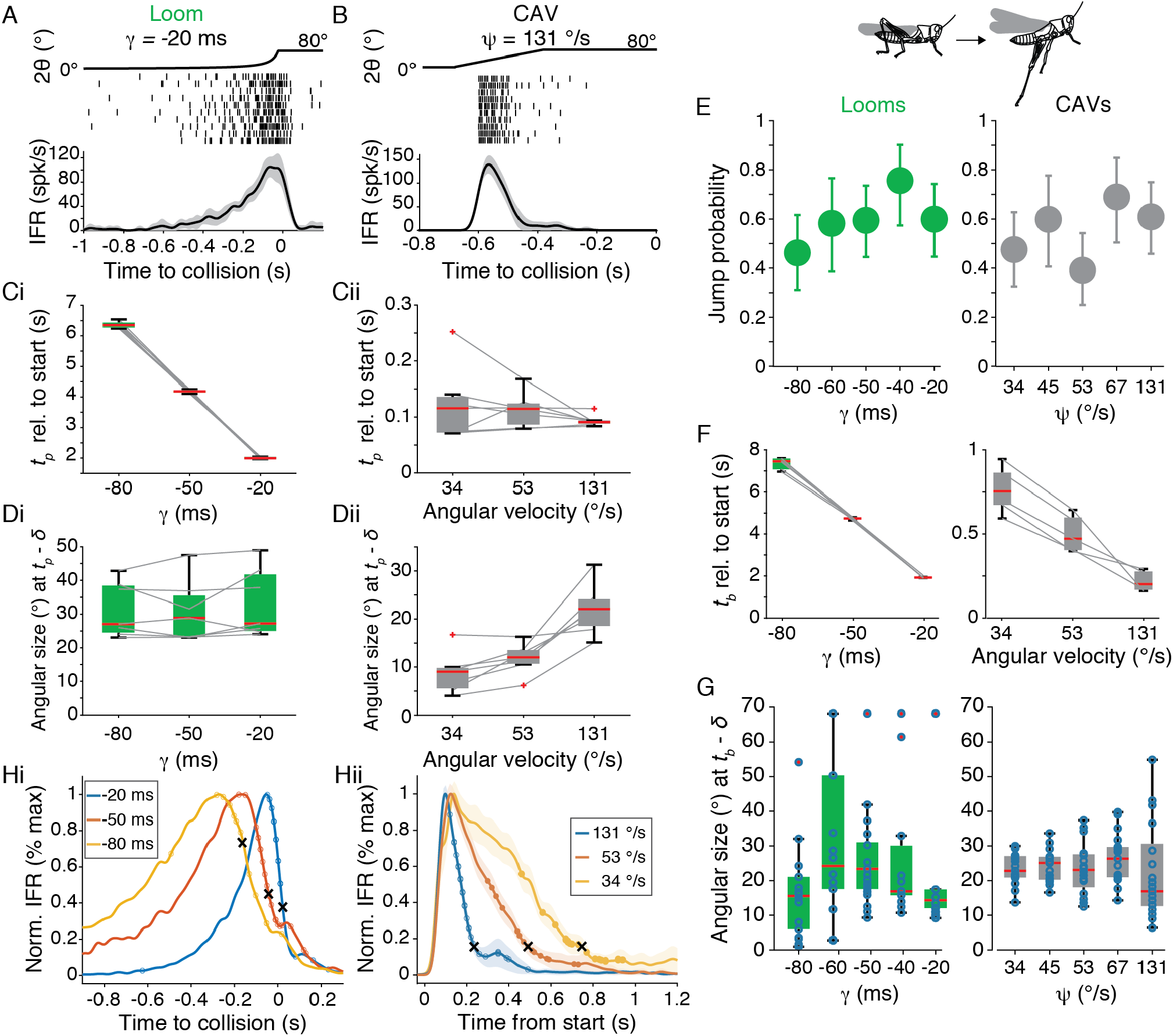
Comparison of responses to stimuli simulating constant approach and constant angular velocity. A) Representative neural responses in a grasshopper to a looming stimulus that simulates an object with constant approach velocity. *Top*, time course of stimulus expansion. *Middle*, spike rasters of DCMD activity, each line is the response to a separate trial. Bottom, instantaneous firing rates (IFR) of the DCMD response presented as mean ± sd. B) Responses of the same animal to stimuli expanding with constant angular velocities (CAVs). Data presented as in (A). C) The time from the start of expansion to the peak response differs for looming stimuli of different expansion rates (Ci, p_KW_ = 1.3•10^−4^, N = 7), but constant angular velocity expansion produces peak response delays unchanged by expansion rate (Cii, p_KW_ = 0.61, N = 7). D) Looming stimuli produced a peak response at a fixed angular size (modulo a time delay, *δ*; Di, p_KW_ = 0.71, N = 7). However, the threshold angular size in response to CAVs depends on angular velocity (Dii, p_KW_ = 0.001, N = 7). E) Grasshoppers jumped in response to both CAVs and looms with equal probability. F) The jump time from the start of expansion was not constant for looms (p_KW_ = 7.5•10^−5^, N = 6) or CAVs (p_KW_ = 3.5•10^−4^, N = 6). G) The stimulus threshold size of the jump was consistent across both CAVs and looms; *δ* = 76 ms. H) Jumps occurred after the neural responses started to decay, as may be seen by comparing the average normalized IFRs in response to looms (i) and CAVs (ii) with the jump times. Individual trial and average jump times are marked with circles and x, respectively.

The same stimuli were presented to freely moving grasshoppers which jumped equally to both looms and CAVs (Fig. 8E). The timing of the behavioral response matched that of the neural response for looms but not for CAVs (Fig. 8F). The escape jumps from CAVs did not come a fixed delay after the start of expansion, but instead after the same threshold angle as for looms (Fig. 8G). Estimating best *η* fits for grasshopper escape times for looms and CAVs yielded no difference in *δ* (p_ASL_ = 0.41) or threshold angle (p_ASL_ = 0.30). These data show that while the grasshopper’s neural response timing to CAVs is better predicted by the *η* model, the behavioral timing is better described by the *κ* model.

Previous work on the sensory-motor transformations leading to escape jumps for looms in grasshoppers has suggested that the decrease in firing following the DCMD peak rate is key in triggering the jump, presumably because it results in decreased flexor excitation (Fotowat & Gabbiani, 2011). We therefore examined averaged neural response profiles to see how they related to the timing of jump escape. As expected, jumps in response to looms occurred on average during the decaying phase of the DCMD response (Fig. 8Hi). Although neural responses to CAVs peaked at the same time, their decrease from peak slowed with decreasing stimulus angular velocity (Fig. 8Hii). The variable rate of decay resulted in an average behavioral response timing occurring at a similar fractional decay from the peak firing rate (∼ 20% on average; Fig. 8Hii).

## Discussion

We used two new classes of laboratory stimuli to study the responses of collision-detecting neurons and escape behaviors in grasshoppers and goldfish. The first class, NZAs, is built on classical looming stimuli by adding constant acceleration or deceleration to interpolate between different initial and final *γ* values. NZAs were inspired by the observation that predator approach trajectories accelerating towards their prey lead to less successful escape behaviors of freely behaving vinegar flies, and vice-versa for deceleration (von Reyn *et al*., 2014). Our experiments establish a similar result in a controlled setting for grasshoppers as accelerating stimuli produce more escapes after the projected collision time (Fig. 4). In contrast, the escape behavior of goldfish shows a considerably reduced sensitivity to acceleration and deceleration, but all escapes occurred before projected collision.

The second stimulus class, CAVs, borrows from stimuli used to elicit escape behaviors in rodents. Those stimuli expand with constant angular velocity but typically span a small angular range (e.g., 2-20º at the animal’s retina) and are presented in short succession 5-10 times to elicit rodent escape behavior (Yilmaz & Meister, 2013; Wei *et al*., 2015; Salay *et al*., 2018). Here, CAVs were presented only once and expanded to cover a large fraction of the visual field like classical looming stimuli (> 80º). In grasshoppers, the use of CAVs confirmed the dichotomy between the LGMD/DCMD neural responses and behavioral escape jumps, with the former better described by the *η* model and the latter by the *κ* model.

The timing of the neural responses to CAVs was a fixed delay, *δ*, after stimulus start as predicted by the *η* model, but the *δ* was ∼100 ms unlike that of looms which is ∼25 ms (Fig. 8Cii). The neural delay between stimuli and LGMD activation is stimulus dependent, with larger and faster stimuli producing shorter delays (Jones & Gabbiani, 2010; <u>unpublished data</u>). For looms and NZAs where the peak response occurs near the time of collision, this peak occurs when the stimulus is large and has a high angular velocity, producing shorter delays of ∼25 ms. For small stimuli the delay to LGMD activation can be >100 ms due to delays in the activation of the presynaptic network (Jones & Gabbiani, 2010). This is the most likely reason for increased delay of the peak responses to CAVs compared to those of looms and NZAs.

In grasshoppers and goldfish, two different models, *η* and *κ* respectively, have been proposed to describe the responses of the LGMD/DCMD neurons and the Mauthner cell to looming stimuli (Gabbiani *et al*., 1999; Preuss *et al*., 2006). These models are difficult to distinguish, though, because they both predict that the time of peak neuronal responses should occur modulo a time delay, *δ*, at a constant angular threshold size in response to looms, independent of their parameter *γ*. One difference between the models is that only the *η* model predicts a constant LGMD/DCMD number of spikes, independent of *γ*. In contrast, only the *κ* model predicts a constant peak M-cell membrane potential (sec. 5, Preuss et al 2006; Gabbiani *et al.*, 2022). Both predictions are in reasonable agreement with experimental observations but represent weak constraints distinguishing the models since they could be implemented in either model through a rescaling of the output by addition of a static non-linearity (sec. 5, Gabbiani *et al.*, 2022). In contrast, the *η* and *κ* models predict different peak response times for NZAs and CAVs, thus opening the possibility of distinguishing the two models through their main predictions: the timing of peak response and the angular threshold size at peak time. Besides distinguishing the two models, NZAs allowed us to investigate the influence of stimulus acceleration on the responses of the looming sensitive LGMD/DCMD neurons in grasshoppers, as well as behavioral escape responses in grasshoppers and goldfish.

Interestingly, a different phenomenological model for the membrane potential of the *Drosophila* giant fiber (GF) in response to looming stimuli has been developed (von Reyn *et al*., 2017; Ache *et al*., 2019). By firing a single spike, this neuron, similar to the Mauthner cell, triggers a fast, short latency jump and subsequent flight escape thought to be of last resort (Card & Dickinson, 2008; Fotowat *et al*., 2009; von Reyn *et al*., 2014). In contrast to the *η* and *κ* models that multiply angular size or speed with a negative exponential of size, the GF model sums four non-linear terms representing the inputs to the GF of four classes of neurons, two excitatory and two inhibitory. Just as the *η* and *κ* models, the GF model peak membrane potential output during looming is tuned to a fixed angular size irrespective of *γ* (Ache *et al*., 2019). But for NZAs and CAVs it behaves more like the *κ* model rather than the *η* model (Figs. 5 and 6, Gabbiani *et al.,* 2022).

In grasshoppers, the neural responses of the LGMD/DCMD neurons agreed with several predictions of the *η* model. Specifically, peak responses to accelerating NZAs occurred later than for equivalent constant speed stimuli while peak responses to decelerating NZAs occurred earlier. Further, the angular threshold size at the peak response time was not constant for accelerating and decelerating stimuli. Yet, the predicted change in angular threshold size at the time of peak firing rate when switching from looms to NZAs was not that observed experimentally. Further, grasshopper escape behavior to NZAs was better described by the κ model, suggesting an additional transform between the descending sensory neural activity and the motor output triggering escape jumps. In grasshoppers, we further tested neural and behavioral responses to CAVs, confirming the results obtained with NZAs.

In goldfish, escape response times to NZAs were more variable than in grasshoppers. This may be expected since multiple pathways drive goldfish escape behavior, in contrast to grasshoppers (Fotowat *et al*., 2011; Domenici & Hale, 2019). The angular threshold size preceding the escape behavior remained relatively constant, irrespective of the accelerating or decelerating profile of the NZAs. This experimental result was consistent with the predictions of the κ model. However, the timing of escape responses showed little sensitivity to acceleration or deceleration, in contrast to κ model predictions. Yet, an increased sensitivity to NZAs could be detected for behavioral reaction times when restricting the analysis to fast bends, likely initiated by the Mauthner cell in response to faster approaches.

Thus, in both species the timing of behavioral responses was more variable and less well predicted by the *η* and *κ* models, respectively, than those of the LGMD/DCMD neurons and the Mauthner cell, as proxied by fast bends. Part of the increased variability in response timing is due to experimental variability in the exact positioning of the freely moving animals, changing the stimulus angular expansion slightly between trials. Additionally there is variability in the behavioral response threshold angle, escape trajectory and escape strategy, that might prevent predators from anticipating their prey’s escape responses (Domenici *et al*., 2011; Bateman & Fleming, 2014).

Evaluating different looming detection computations raises the question of what predator detection implementations would be most effective for survival. The current lack of data on predator approach trajectories means this question cannot yet be answered precisely, but some general points can be inferred. All animals need to detect collision early enough to account both for the time required to execute the escape and the delays of neural processing. Many prey animals, including grasshoppers, are most visible to predators when fleeing and at least one bird species tries to visually evoke escapes to improve predation (Jabłoński & Strausfeld, 2001). So, prey may want to delay escape until it is necessary.

Using a constant threshold angular size to trigger escape is advantageous if the threshold angle is small enough to give time to escape and large enough to avoid most false alarms. The shift in escape timing predicted by a strict implementation of the *η* model may be maladaptive, in that it would cause escapes at smaller angles for decelerating approaches that require less time to flee from, and at larger angles for accelerating approaches that provide less time to escape (Fig. 2H). The more effective escape strategy would be to adopt a smaller threshold angle for accelerating stimuli, which is not accomplished by a strict implementation of either *η, κ*, or GF models.

Grasshoppers, unexpectedly, exhibited such a decrease in threshold angular size for accelerating stimuli (Fig. 7B), suggesting there may be additional visual processing to escape accelerating predators more effectively. Despite this, though, most responses to stimuli with high acceleration occurred after the projected collision time (Fig. 4D). The speed of the Mauthner cell triggered escape response in goldfish might make such a shift in threshold angle for accelerating stimuli unnecessary as all recorded responses were before projected collision.

In summary, responses to accelerating stimuli support the *η* and *κ* models for the LGMD/DCMD neurons and the Mauthner cell, respectively, while also showing their limitations. The behavioral response of grasshoppers was better described by the *κ* model, though, revealing that sensory information is transformed as it interfaces with the motor system driving escape jumps. As a result, collision avoidance in grasshopper and goldfish converge towards a common escape strategy relying on angular threshold size irrespective of stimulus properties, as reflected in the κ model. The same stimuli could help disambiguate competing models of looming responses in other species and offer further insights in the neural computations underlying escape behaviors. As more is learned about the approach trajectories of predators, understanding the underlying computations serving collision-avoidance will also provide insight into the neural adaptations resulting from predator-prey interactions.

## Additional Information

### Data availability statement

All data presented in the manuscript and the Matlab code to generate the figures is available on Mendeley Data (repository DOI).

### Author contributions

FG, RD and TP designed the work. RBD, TC-M and ME carried out the experiments and analyzed the data. RD, FG and TP wrote the manuscript, with additional input from TC-M and ME.

All authors approved the final version of the manuscript. All persons designated as authors qualify for authorship, and all those who qualify for authorship are listed.

## Acknowledgments

Thanks to Ms. Gona Kordestani for technical help with grasshopper experiments. Thanks to the BCM Bioengineering Core for aid in experimental design (supported through an NEI Core Grant for Vision Research, EY-002520-37).

